# CBD lengthens sleep, shortens ripples and leads to intact simple but worse cumulative memory

**DOI:** 10.1101/2023.02.28.530388

**Authors:** Anumita Samanta, Adrian Aleman-Zapata, Kopal Agarwal, Pelin Özsezer, Alejandra Alonso, Jacqueline van der Meij, Abdelrahman Rayan, Irene Navarro-Lobato, Lisa Genzel

## Abstract

Cannabidiol (CBD) is on the rise as over-the-counter medication to treat sleep disturbances, anxiety, pain and epilepsy due to its action on the excitatory/inhibitory balance in the brain. However, it remains unclear if CBD also leads to adverse effects via changes of sleep macro- and microarchitecture. To investigate the effect of CBD on sleep and sleep-related memory consolidation, we performed two experiments using the Object Space Task testing both simple and cumulative memory in rats. We show that oral CBD administration extended the sleep period but changed the properties of NonREM sleep oscillations (delta, spindle, ripples). Specifically, CBD also led to less long (>100ms) ripples and consequently worse cumulative memory consolidation. In contrast, simple memories were not affected. In sum, we can confirm the beneficial effect of CBD on sleep, however, this comes with changes in NonREM oscillations that negatively impact memory consolidation.

## Introduction

Cannabidiol (CBD) has become a popular remedy for many ailments and shows promising effects treating chronic pain, anxiety and sleep disturbances [1, 2]. In patients with neuropsychiatric disorders, CBD administration improved sleep quality in more than half of the patients up to a month after administration and alleviated anxiety symptoms [3]. Furthermore, due to the action on inhibition in the brain, CBD is potentially a novel therapeutic agent for restoring excitation/inhibition balance in models of epilepsy [4] and autism, which stem from dysregulation of inhibitory networks [5]. Interestingly, the effect of CBD on sleep is dose-dependent with high doses increasing sleep duration [6] and the opposite effect is observed with low doses [7]. Despite the rising use of CBD by various patients as an over-the-counter remedy, very little is known about how CBD affects the microstructure and functions of sleep, which could lead to side-effects as seen with other sleep-modulating medications [8–10].

Sleep is thought to play a critical role in the consolidation of memories, both for simple memories – such as the association of two items – as well as the more complex extraction of regularities across different experiences. NonRapid-Eye-Movement (NonREM) sleep has been proposed as a critical stage in offline memory processing [11, 12]. The reactivation of cells active during encoding, also referred to as *neural reactivation*, has been shown to occur during quiet wake and NonREM sleep and is potentially crucial to memory consolidation [13–16]. These reactivation events occur during hippocampal high frequency burst oscillations (100-250 Hz) referred to as ripples [17]. To be able to consolidate memories during sleep, a coordinated action between the hippocampus and the neocortex around the ripples is needed. Two major neocortical events – delta waves (0.5-4 Hz) and spindles (9-20 Hz) – can occur in close coordination with the ripple, and this crosstalk between brain regions is crucial for offline memory processing [18–21]. The characteristics and coordination of these microstructural sleep events are influenced by the depth of sleep as well as most sleep-inducing medications [9, 10]; thus, there are likely effects of CBD on these oscillations that could negatively impact their function in memory consolidation.

CBD acts as an antagonist of the CB1-receptors [22–24], which are present in cholecystokinin (CCK) and parvalbumin (PV) positive interneurons in the hippocampus [25]. CB1-receptors are found in the presynaptic axon terminals of the GABAergic interneurons, most prominently in the CA1-CA3 subfields [26, 27]. They act like retrograde messengers, wherein their activation in the presynaptic targets is triggered by depolarization in the post-synaptic targets [25]. CB1-receptors potentially regulate the suppression of GABA-mediated transmission following the depolarization of hippocampal pyramidal neurons [28, 29], which is a key player in the occurrence of ripples. The ripple component is initiated by the activation of pyramidal neurons in the CA1, in response to input from CA3; however, the oscillation is maintained by interneuron activity [30]. Thus, by changing the E/I balance, CBD is likely to have an impact on ripples and on memory consolidation.

To test the effect of CBD on sleep and sleep-related memory consolidation, we performed two experiments using the Object Space Task [31]. This task enables measuring both simple and cumulative memory, where the former should be more resistant to manipulation, whereas the latter should be more sensitive to manipulation effects. In the first experiment, we tested the effect of oral CBD administration – mimicking human CBD intake – before learning and consolidation on memory expression the next day. For the second experiment, we conducted in fewer animals recordings of natural sleep and sleep oscillations after learning with electrophysiological implants targeting the hippocampus and the cortex to elucidate how CBD affects the sleep macro- and microstructure. Oral administration of CBD extended sleep at the end of the sleep period. Furthermore, it changed sleep microstructure – leading to less long ripples – that resulted in worse cumulative but intact simple memories tested the following day.

## Methods and Materials

### Study design

A total of 48 rats were used in this study: 44 rats were used for behavioral experiments and four rats were used for chronic electrophysiological recordings. All rats were first extensively handled for multiple days until they experienced minimal stress while working with the experimenters (see handling videos on www.genzellab.com). The first batch of 44 rats was first habituated to the oral feeding regime and the behavioral training box for a week. In the following week, they were orally administered with >98% trans-cannabidiol (CBD) or vehicle (VEH) (counterbalanced across animals) and trained in the Stable and Overlapping condition of the Object Space task (all conditions run cross-over within each animal) in smaller sub batches of 14-16 animals (maximum 8 in one day).

The 4 implanted rats underwent surgery for wire drive implantation. After surgery, they were allowed to recover for up to 2-3 weeks, during which the wire arrays were slowly lowered to reach the pyramidal layer in the CA1 region of the hippocampus and the rats were habituated to the behavior training box and the oral feeding regime. Upon reaching the target, we started the behavioral experiments. Similar to the behavior batch, these rats were orally administered with either CBD or VEH and then trained in the Object Space Task (all conditions cross-over within subject). Brain activity was recorded during exploration and rest periods.

### Animals

Six-to eight-week-old male Lister Hooded rats weighing between 250-300g at the start of the experiment (Charles Rivers, Germany) were used in this study. They were pair-housed in conventional Eurostandard type IV cages (Techniplastic, UK) in a temperature-controlled (20 + 2 ºC) room following a 12h light/dark cycle with water and food provided *ad libitum*. The first batch of 44 rats was kept under these conditions until the end of the experiment. The batch of chronic rats were single-housed after wire drive implantation and kept in these conditions until the end of the experiment. A total of 48 rats were used in this experiment: 44 in the behavioral experiment (3 sub batches of n=16, 14 and 14), and four for chronic electrophysiological recordings (two more served as implant-pilots but were not included in the study). The behavioral experiments and electrophysiological recordings were performed during the light period (between 9:00-18:00). All animal procedures were approved by the Central Commissie Dierproeven (CCD) and conducted according to the Experiments on Animals Act (protocol codes, 2016-014-024 and 2020-0020-006).

### CBD

(-)-trans-Cannabidiol (CBD, > 98%) was obtained (https://www.cbdepot.eu/products/cannabidiolum-gmp) for all experiments. Rats were treated with either CBD (120 mg/kg in 300 μl flavored olive oil, p.o.) or vehicle (300 μl flavored olive oil, p.o.). Different flavoring agents, namely, vanilla, cinnamon, star anise and clove, were used to make multiple flavors of olive oil, which were then used to mask the taste of CBD, so that the rats would always be naïve to what they were fed. The use of all flavors was counterbalanced, such that each rat received all flavors for both CBD and VEH. CBD solution was always freshly prepared in one of the flavored oils prior to oral administration. In order to prepare the solution, first the amount of CBD to be administered was determined based on the rat’s weight and the compound was weighed accordingly. The flavored oil was heated up to a temperature of 50-60°C and then the CBD compound was slowly dissolved into it. This process would take a few minutes until a clear solution was obtained, which is indistinguishable from vehicle oil. Individual syringes with 300 μl CBD or VEH were prepared for each rat. The experimenter who performed the oral administration was blinded to the treatment each rat received, and the drug was always administered an hour before the start of behavioral training. Further, the experimenters performing the behavioral training and electrophysiological recordings were blinded to the treatment of the animal as well. Previous studies have shown oral administration of CBD to be effective in crossing the blood brain barrier and plasma concentrations are shown to reach their maximum around 4-6 hours after oral administration, which is optimum time window to examine its effect on memory consolidation [32, 33].

### Behavior - Object Space Task

The Object Space Task was used in this study to assess the effects of CBD on simple and semantic-like memories [31]. This task is based on the tendency of rats to explore novel object locations in an open field arena across multiple trials. The animals were extensively handled by the experimenters in the first week to minimize their stress levels during the interactions. Next, they were habituated to the training box (75 × 75 cm) for five sessions over five days. On the first day, they spent 30 min in the training box with their cage mates. On the second and third days, each rat individually spent 10 mins exploring the training box. For the final 2 days, two objects (made from DUPLO blocks, not used in main experiment) were placed in the center of the box and the rats were allowed to explore for 10 mins each day.

A total of 44 rats were trained in the Stable and Overlapping conditions of the task and ran in smaller sub batches of 8. The condition sequence and object locations were counterbalanced across all rats and the experimenter was blinded to the conditions and drug treatment. We followed a within-subject design, wherein every rat performed all conditions of the task with CBD and VEH.

For rats, running one round of one condition took two days – a training day and a test day 24 hours later. The rats were first orally administered CBD or VEH and then started with behavioral training an hour after drug administration. The training day consisted of five trials, 5 mins each with an inter-trial interval of ~45-50 mins. During each trial, the rat was placed in the training box to explore a pair of identical objects at fixed locations. The object pairs were changed from trial to trial. Additionally, to prepare for one session of training and test, the training box was equipped with 2D and 3D cues to create a unique spatial environment for the rats to orient themselves. This box setup remained the same for one session of training and test, and was changed as much as possible for the next session, in order to create a distinctly different environment each time. In the Stable condition, the positions of the objects remained the same across all trials. 24 hours later, during the test trial, one object was moved to a different position. This condition allowed us to assess simple memory in the rats. For training in the Overlapping condition, one position of the object remained constant across all trials whereas the other kept changing. However, the object positions in the final trial of the training day and the test day, 24 hours later, were kept the same. If the rat formed a cumulative memory of one object position constantly moving across multiple trials on the training day, he would spend more time exploring that position. This would let us assess semantic like memory in the rats. Implanted animals additionally had Home Cage days, where they would have the same daily regime as on a training day but instead of being placed in the training box for the 5 min training periods, the animal was kept awake in the home cage. Therefore, these days are non-learning controls that are controlled for general wake activity/sleep disturbances of training and time of day.

All behavior sessions were recorded using a webcam placed above the training box. The object exploration times were scored online in real time during the trials using a scorer program developed in the lab for training and scoring (https://github.com/NeuroNetMem/score). Whenever a rat would sniff or climb on or interact with an object, it was scored as an exploring behavior. Obsessive chewing or biting of an object was not considered as an exploratory behavior, and extra care was taken to avoid using objects that would trigger these behaviors. The scoring data from the behavior sessions was saved in an Excel sheet and was used for further analyses.

### Bilateral wire drive implants

A customized lightweight wire drive implant was manufactured in collaboration with 3Dneuro (https://www.3dneuro.com/) to implant bilateral wire arrays in the prelimbic cortex and the CA1 region of the hippocampus. Customized individual wire arrays (Science Products, catalog no. NC7620F) were first built for each brain region. The wire arrays for the prelimbic cortex consisted of four wires of the same length glued to each other; for the hippocampus, the array consisted of four wires of different lengths with a 70° angle between them to enable recording from the pyramidal layer as well as from below and above the layer. These arrays were then glued to polyamide tubes which were glued to the 3D printed drive body. The drive body was designed according to the Rat Brain Atlas in Stereotaxic Coordinates [34] to ensure the correct placement of arrays in the regions of interest. Additionally, a shuttle and screw were included in the drive body design for the hippocampal arrays, so that they could be turned down after implantation to reach the target layer. Finally, the drive body was designed to enable recording of each region from both hemispheres. Once all the arrays were loaded into the drive body, their free ends were connected to a customized 32 channel electrode interface board (EIB) using gold pins (Neuralynx). A single NPD dual-row 32 contact omnetics connector was attached to the EIB to connect the headstage for later recordings. The hippocampal wire arrays were flushed with the polyamide tubes before implantation. The bottom of the drive was deepened in 70% ethanol for at least 2 hours before implantion into the brain.

### Drive implant surgery

Shortly before the start of surgery, all rats received a subcutaneous (sc) injection of carprofen (5mg/kg) to serve as an analgesic. After placing the rats into the stereotaxis, they received a subcutaneous injection of a mixture of 4 mg/kg lidocaine and 1 mg/kg bupivacaine in a 0.9% NaCl physiological serum locally at the skin surface above the skull as a local analgesic. Finally, they also received a 2 ml of 0.9% NaCl physiological serum subcutaneous injection at the start and end of surgery. Since this was a recovery surgery, utmost care was taken to maintain the most sterile conditions possible and the entire surgery was performed under isoflurane inhalation anesthesia. Additionally, the rats were administered a 10 mg/kg subcutaneous injection of Baytril antibiotic at the beginning of surgery to prevent postsurgical infections. Two pairs of holes were drilled bilaterally to reach the following targets – prelimbic cortex (AP +3.5mm and ML +-0.5mm) and hippocampus (AP −3.8mm and ML +- 2mm). Additionally, a ground screw (M1×3mm) was placed on the right hemisphere of the cerebellum. Three more M1×3mm supporting screws were placed and bound to the skull using Superbond C&B dental cement (Sun Medical, Japan). Upon drilling all holes above the target regions, the dura mater was carefully removed, exposing the surface of the brain. Finally, the wire-drive was carefully positioned on the brain’s surface, such that the wire arrays fit well in the drilled holes. On finalizing the position, the drive was attached to the skull and screws by simplex rapid dental cement (Kemdent, UK). The wire arrays for HPC were slowly lowered (~1.5mm DV from brain surface) to a layer close to the pyramidal layer in the CA1 region. The pyramidal layer was reached progressively in the subsequent days during signal checks in the rats’ recovery period. With one out of the 4 rats, we were not able to reach the pyramidal layer. This rat was thus included only for analysis of cortical events and sleep architecture analyses.

### In vivo electrophysiological recordings

The animals were closely monitored for a week following the drive implant surgery to ensure they had a good recovery. Their weights were monitored daily during this period to ensure a stable growth curve. The rats were allowed to recover for a couple of weeks following surgery before starting with recordings. Prior to the surgery, the rats were habituated to the sleep boxes, chocolate treats and the oral feeding regime, to minimize any stress post-surgery. During the recovery period, they were re-introduced to the sleeping boxes and their local field potential (LFP) activity was monitored during wake and rest periods to assess the placement of the wire arrays. The HPC arrays were further lowered in small steps during this period and the correct placement was confirmed with the LFP activity. Finally, after a week of recovery, the rats were also habituated to the Object Space box according to the habituation protocol described above.

Once all arrays were in the target regions, recordings were started while the rats were being trained in the Object Space task and during the in between rest periods. The feeding and training protocol that was followed here was similar to the one described above. Each rat had a session per experimental condition (Homecage, Stable, Overlapping) with prior administration of CBD or VEH. In the end, every rat ran each session, once with CBD and another with VEH. The sequence of conditions was counterbalanced for every rat and the experimenter scoring the behavior was blinded to which drug treatment the rat was assigned to.

The LFP and accelerometer activities detected by the channels were amplified, filtered and digitized through a 32 channel headstage (IntanTechnology) connected through an Intan cable and a commutator into the Open Ephys acquisition box [35]. The signal was visualized using the open source Open Ephys GUI at a sampling rate of 1 Hz – 30 kHz.

#### Electrolytic lesions

After the final recording session, the animals received electrolytic lesions under isoflurane inhalation anesthesia 48h before perfusion to identify the electrode tips placement. To this end, a current of 8 μA for 10 s was applied in two wires per array using a stimulator.

### Histology

#### Brain processing

Rats were sacrificed via transcardial perfusion 48 hours after the electrolytic lesions. They were overdosed with 150 mg/kg sodium pentobarbitol ip prior to perfusion. They were perfused first with 100 ml of 0.1 M phosphate-buffered saline pH 7.4 (PBS) followed by 250 ml of 4% paraformaldehyde (PFA) made in 0.1 M PBS. After extracting the brains, they were kept overnight in PFA at 4°C. The brains were rinsed in 0.1 M PBS the next day (3 × 10 mins) and then kept in a solution of 30% sucrose, 0.02% NaN3 in PBS for cryoprotection. Once the brains sank to the bottom of the vial, they were frozen in dry ice and stored for the long term in a −80°C freezer. For further processing, the brains were sectioned in a cryostat (SLEE medical, Germany) and 50 micron coronal sections of target regions were obtained and collected in 48-well plates containing 0.02 % NaN_3_ in PBS and stored at 4°C.

#### Nissl staining

Coronal sections containing the prelimbic cortex (PFC) and hippocampus (HPC) were sequentially mounted (in increasing AP coordinates) on gelatin-coated slides and incubated overnight at 37°C. The following day, the slides were further processed for Nissl staining. Slides were first hydrated in 0.1 M PBS and then in Milli Q water for 20 min each. Next, the slides were stained in 0.7% acetate Cresyl Violet for 20 min and dehydrated in an increasing ethanol gradient (water for 3 min, 70% ethanol for 20s, 96% ethanol + acetic acid for 45s, 100% ethanol for 5 min). Finally, in the last step, the slides were immersed in xylene for 15 min and then then mounted with DePeX mounting medium and covered with coverslip. Lesions from wire arrays were then observed under a bright field microscope (LEICA DM IRE2) and images were taken in 5X and 10X magnification.

### Behavioral Data Analyses

The amount of time the rats spent exploring objects was scored in real time which would then get saved in an excel sheet. The total exploration time was calculated as the sum of time spent exploring both objects. Further on the discrimination index (DI) was calculated by subtracting the amount of time exploring the familiar location from the novel location and dividing by the total exploration time. DI > 0 indicated a preference for the novel object location, which was used as a prime measure for memory performance. DI = 0 indicated no preference for either location and finally DI < 0 indicated a preference for the stable object location. Data was analyzed with rmANOVA (within subject factors treatment, condition).

### Local field potential analysis

Here, we hereby describe the methods followed to process and analyze the local field potential data acquired during chronic recordings. The scripts used to implement this can be found in the following github repository https://github.com/genzellab/cbd. All the recorded data is in the Donders repository (di.dcn.DAC_626830_0008_841). A downsampled version of the LFP data can be found at https://osf.io/fwshb/.

#### Scoring of sleep

Recordings from the rest periods across all study days were further classified into different sleep/wake states using a sleep scoring GUI called TheStateEditor in MATLAB developed by Dr. Andres Grosmark [36]. As the first step, one PFC and HPC channel and three accelerometer channels were selected per animal as input into the scorer interface. One could then visualize the frequency spectrograms and bandpass filtered LFP signal per brain region and the motion spectrogram during the recording period. Using this information, an experienced researcher scored the recording data in increments of 1 second epochs into one of the following states – wakefulness; NonREM; Intermediate; REM. The researcher scoring the datasets was blinded to condition and treatment for that respective study day. Absence of movements in certain periods in the motion spectrogram further aided in discriminating between sleep and wake periods. Desynchronized activity in PFC and HPC accompanied with movement in the motion spectrogram was classified as a wake state. NonREM sleep was classified when slow oscillations (0.5-4 Hz) were detected in the PFC channel accompanied by lack of movement in the motion spectrogram. REM sleep was classified when we detected desynchronized activity in the PFC channel and dominant periodic theta activity (6-8 Hz) in the HPC channel accompanied by lack of movement in the motion spectrogram. Intermediate sleep was defined as short transition periods between NonREM and REM states which were characterized by irregular theta activity in the HPC channel and high frequency desynchronized activity in the PFC channel accompanied by lack of movement in the motion spectrogram.

#### Sleep architecture analysis

Based on the manual scoring of the states, we computed the total sleep time (TST), wake, and total time spent in different sleep states (NonREM, REM and Intermediate) per session in MATLAB. Averages for these variables were further calculated across all sessions per rat and the mean and SEM was computed per treatment. Additionally, % of TST spent in NonREM, REM and intermediate was calculated in 45 min time bins for the 3-hour rest periods after the final trial. Furthermore, the distribution of bout duration per sleep stage was calculated for every session across different treatment groups. A bout was defined as a continuous period spent in a specific sleep state and only bouts longer than 4 seconds were considered for analysis. The state of switching from one sleep stage to another was defined as transitions. Using MATLAB, we computed the number of bouts per sleep stage, the duration of the bouts, and transitions between sleep stages, for example, number of NonREM-REM and REM-NonREM transitions. This was computed per session for each rat and later averaged for CBD and VEH treatment groups across rats. The MATLAB scripts for this analysis can be found at https://github.com/genzellab/sleep_architecture.

#### Preprocessing of recordings

A prelimbic cortex channel with large slow oscillations during NonREM sleep was selected per rat. A hippocampal channel from an electrode placed in the pyramidal layer was selected based on the quality of ripples visually detected on the channel. The channels were downsampled from 30 kHz to 2.5 kHz by first employing a 3rd order zero-phase Butterworth low pass filter with a cutoff of 1.25 kHz to prevent aliasing of the signal. Artifacts observed during NonREM bouts were subject to a blanking method, which consisted of applying an amplitude threshold determined visually. Artifacts crossing the threshold were “blanked” by replacing them with the mean value of the NonREM bout. For some types of artifacts, a preceding build-up period of one second was also blanked. The artifact removal was done after the signal had already been bandpass-filtered in the corresponding frequency range used for the detection of ripples, spindles or delta events. This prevented artifacts due to discontinuities.

#### Detection of hippocampal ripples

The downsampled channels (2.5kHz) of the hippocampal pyramidal layer were loaded into the MATLAB workspace and their NonREM bouts were extracted. Using a 3^rd^ order zero-phase Butterworth bandpass filter, the epochs of HPC signal were filtered to a frequency range of 100-300 Hz. A custom MATLAB function was used for detecting the start, peak and end of the ripples by thresholding voltage peaks which lasted a minimum duration of 30 ms above the threshold. The start and end of the ripple were determined as half the value of the detection threshold. A closeness threshold of 50 ms was used to count ripples occurring within the proximity of each other as a single event. The detection threshold was determined by computing the standard deviations of concatenated NonREM bouts individually for pre and post sleep trials in a study day. These standard deviations were multiplied by a factor of 5 and an average was calculated to find a single detection threshold per study day. An offset of 5 μV was added to the threshold to reduce false positives. Only pre- and post-trial sleep periods with more than 3 minutes of NonREM were included. This process was repeated for all study days pertaining to all rats in both treatment groups. The detections were then grouped as long (>100ms) or short (≤100ms) ripples based on their duration, in accordance with Fernandez-Ruiz et al. 2019.

#### Detection of cortical spindles and delta waves

A downsampled prelimbic cortex channel (2.5kHz) with large slow oscillations was selected and using a 3^rd^ order zero-phase Butterworth filter the signal was filtered to 9-20 Hz for detecting spindles and to 1-6Hz for detecting delta waves. The NonREM bouts were then extracted from the filtered signal and concatenated. The functions FindSpindles and FindDeltaWaves from the Freely Moving Animal (FMA) toolbox (http://fmatoolbox.sourceforge.net)[37] were modified to adapt the thresholds for optimal detections and were then used to detect the start, peak and end of spindles and delta waves, respectively. The optimal threshold was found for each rat by visually inspecting the detections and modifying the default parameters of the functions when needed. The results were saved as timestamps in seconds with respect to the concatenated NonREM signal. They were then used to find the timestamps with respect to the recorded signal. This process was repeated for pre- and post-trial sleep periods with more than 3 minutes of NonREM in study days pertaining to all rats in both treatment groups.

#### Detection of oscillation sequences

The sequences between ripples, spindles and delta waves were counted in various combinations to study cortico-hippocampal coupling during NonREM sleep as done by Maingret et. al. 2016. The time difference between the peaks of these events was compared to a fixed duration to establish if there was a sequential relationship in the following combinations of oscillations: Delta-Spindle (D-S), Delta-Ripple (D-R), Ripple-Spindle (R-S), Delta-Ripple-Spindle (D-R-S), Delta-Spindle co-occurring with Ripple (D-SwR). For D-S a sequence was considered when the duration between events was between 100-1300 ms, for D-R it was 50-400 ms and for R-S it was 2-1000 ms. To find D-R-S sequences, the results of D-R and R-S were compared to find ripples preceded by a delta wave and followed by a spindle. To find the D-SwR sequences, the results of D-S and spindles co-occurring with ripples (see next subsection)were matched to find spindles preceded by a delta and co-occurring with a ripple. The results were saved as counts of each sequence for each pre- and post-sleep trial. This analysis was then repeated using timestamps of long and short ripples separately.

#### Co-occurrence between ripples and spindles

The co-occurrence between ripples and spindles was computed by comparing the start and end timestamps of both events. To consider co-occurrence between a ripple and a spindle, either one of the following conditions had to be fulfilled: 1) A ripple had to start and end within the duration of the spindle. 2) One of the events had to start or end within the duration of the other. Given that more than one ripple could co-occur with the same spindle, we counted separately ripples co-occurring with spindles and spindles co-occurring with ripples. This analysis was then repeated using timestamps of long and short ripples separately.

#### Slow oscillation phase analysis

The downsampled PFC signal (2.5kHz) was filtered to the 0.5-4 Hz range using a 3^rd^ order Butterworth zero-phase bandpass filter and its Hilbert transform was computed to find the phase angle of slow wave oscillations in the range of 0° to 360°. The peaks of long and short ripples were then used to find the corresponding phase and this process was repeated for all study days pertaining to all rats.

#### Ripple rate at bout start and end

NonREM bouts were isolated and bouts longer than 5 mins were used for further analysis. For these bouts, 10% of their duration from the start and 10% at the end of the bout were taken and compared with the timestamps of long and short ripples to get the count of ripples occurring within these segments. The rates were then calculated by dividing the counts by the duration of the 10% segments. This process was done for all study days pertaining to all rats.

#### Oscillations characteristics

The traces of each detected event (ripples, spindles, delta waves) were extracted using the start and end timestamps obtained from the detectors. The traces of the events were filtered in their corresponding detection frequency band. Characteristics such as the amplitude and mean frequency were calculated for these filtered events using built-in and custom MATLAB functions. Namely, the amplitude of the events was calculated by computing the envelope of the filtered trace using a Hilbert transform. The absolute value of the result was taken and its maximum was found. The mean frequency of the filtered traces was computed using the meanfreq function of MATLAB.

#### Baselines generation for power spectra, and slope analysis of power spectral density

NonREM bouts were extracted from three rats, six study days per rat (18 study days in total) which include presleep and post-trials sleep recordings. The six study days per rat consisted of all three conditions (i.e., Overlapping, Stable, and Home Cage) per treatment (i.e., VEH and CBD). Bouts shorter than 4 seconds were not considered. Finally, 400 NonREM bouts were randomly selected from all NonREM bouts while epochs with a time window of 4 seconds were randomly extracted from each of these NonREM bouts. The extracted windows contained simultaneous activity in PFC and HPC. Baselines were only included for power spectral density (PSD) slope analysis.

#### Power spectra & power spectral density slope analysis

Power spectrum and slope analysis of power spectral density were computed for chronic recordings (i.e., HPC and PFC). More specifically, they were computed for both VEH and CBD treatments. Baselines were randomly selected NonREM periods of 4 seconds. First, the Fieldtrip toolbox (Oostenveld et al., 2011)[38] was used to compute PSD. PSD was computed from 0 to 100 Hz in steps of 0.25 Hz and with a length of 4 seconds. Line noise was removed at 50 Hz using a Notch filter. A Hanning window taper was used with a length of 4 seconds with a 0.25-second overlap. To generate power spectra, the PSD values were computed as logarithms with a base of 10. To calculate the slope and offset, the ‘Fitting Oscillations and One-Over-f (FOOOF)’ Python toolbox was used (Donoghue et al., 2020)[39]. Please note that it is advised to pass in non-logarithmic power values to the FOOOF toolbox as the toolbox handles computing logarithm itself. The minimum peak height threshold was set to 0.05 (units of power – same as the input spectrum), and peak width limits were between 1 Hz and 8 Hz. The outputs were in logarithms with a base of 10. The resulting values of slopes and offsets were analyzed using IBM SPSS Statistics (Version 29)[40]. For slopes and offsets, a two-way ANOVA was conducted to determine the effects of treatment (VEH & CBD) and the ripple type (random NonREM baseline, short & long ripples). Statistical significance was accepted at the p < 0.05 level for simple interactions and simple main effects. Post hoc comparisons using the Tukey HSD test were conducted to investigate interaction effects.

#### Power Spectra of Sleep Stages

The PSD for REM and NonREM was first calculated for six study days per rat (18 study days in total), and one presleep and all five trials of each study day using the Welch method with a window length of one second and 80% overlap. To examine the two components (i.e., slope and offset) of the electrophysiological power spectrum, the Python toolbox ‘Fitting Oscillations, and One-Over-f (FOOOF)’ was used between 0 and 100 Hz.

Slopes and offsets were analyzed using IBM SPSS Statistics (Version 29). A three-way ANOVA was conducted to determine the effects of treatment (VEH & CBD), brain area (HPC & PFC), and sleep stage (REM & NonREM). Statistical significance was accepted at the p < 0.05 level for simple two-way interactions and simple main effects. Post hoc comparisons using the Tukey HSD test were conducted to investigate interaction effects.

## Results

### CBD decreases cumulative memory expression the next day but leaves simple memory intact

Our first aim was to establish a behavioral effect of CBD administration targeting memory consolidation. For this, the Object Space Task (Fig. 1A) was combined with oral administration of CBD, in a similar dose as commonly used in oral administration to humans. CBD and vehicle-control was given 1h before training start (1h after light on, Fig. 1B), such that due to slow uptake in the gastro-intestinal system and crossing of the blood-brain barrier the peak of CBD in the brain occurred after training during the main sleep period [32, 33]. However, some CBD will also be absorbed in the mouth and therefore reach the brain earlier (bypassing liver metabolism). Of note, all experiments were run counterbalanced and blinded. The experimenter performing the behavioral training was not aware of the current drug treatment of the rat.

**Fig.1.**
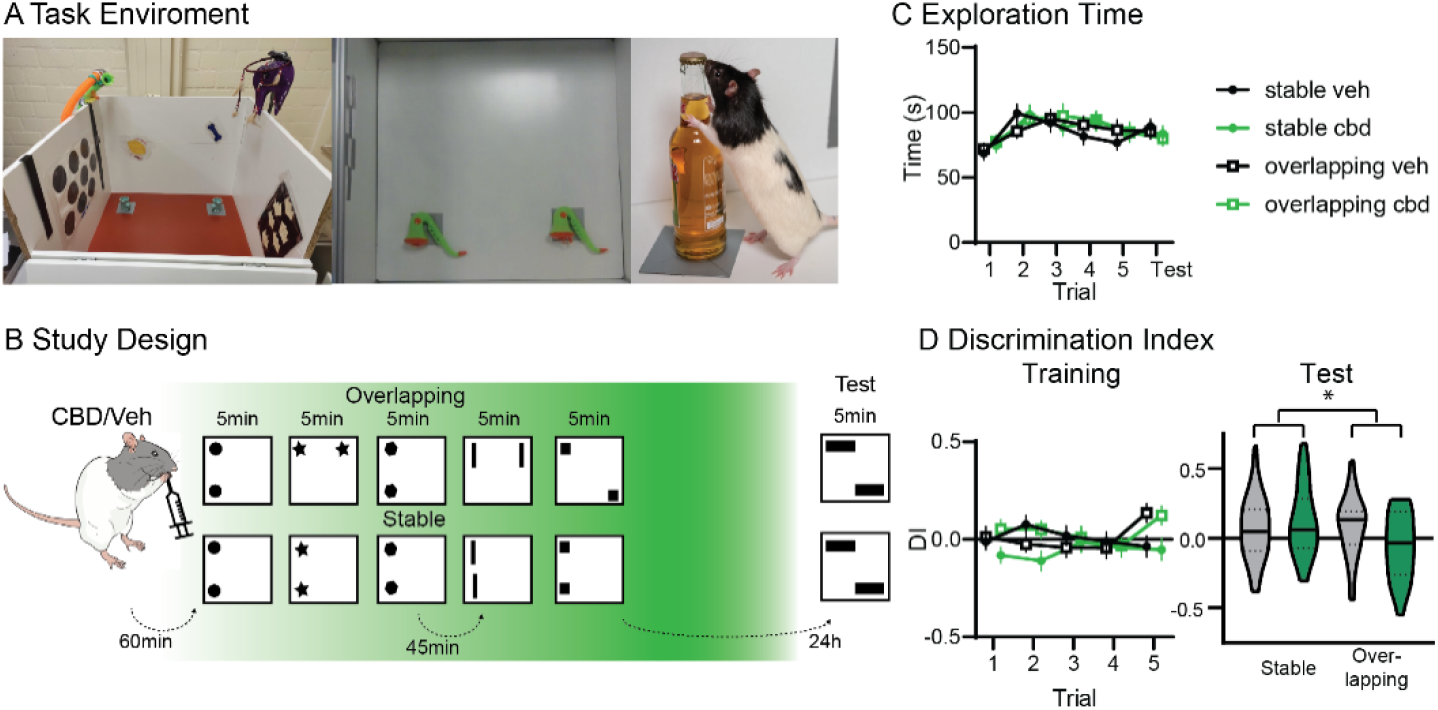
Behavior. A. Shown is the object exploration box (side view, top view and with animal). B. Study Design. Rats received orally either vehicle or CBD (cross-over) and 60min later were trained in the OS task with either Overlapping (one stable, one moving object location) or Stable (two stable object ‘locations) condition (5min trials, inter-trial-interval 45min) and ‘ tested 24h later. C. The exploration time remained stable for both conditions and treatment. D. There was no difference in discrimination index (DI) during training for treatment, but as expected in the fifth trial both showed positive DIs in overlapping. At test, there was a significant condition X treatment interaction (p=0.031). *p<0.05

Rats were trained in either the Stable or Overlapping condition, which test simple and cumulative memory respectively. In Stable, in each training trial identical object pairs (each trial different ones) are presented in the same two locations in the exploration box over 5 training trials. At the test (24h after training), two new objects are again used and presented with one on a usual location but the other one placed in one of the other two corners. Neophilic animals will explore the object in the novel location more, independent of whether they just remembered the last experience or all five training trials (leading to positive Discrimination Index, DI). Thus, the Stable condition tests for simple location memory. In contrast, in Overlapping, already during training only one location remains stable, while the second object will be in one of the three other corners. Now, the test trial has the same configuration as the final training trial. Therefore, only if the rats created a cumulative memory, abstracted over multiple training trials, will they show a positive discrimination index at test. All animals underwent both conditions and both treatments, therefore four rounds of training-test, in a cross-over design. Treatment/condition sequences were counterbalanced across ratss and object locations.

While CBD was administered before training, due to the slow uptake the main effect should be after training in the sleep period. In confirmation, that CBD did not affect encoding behaviors, both exploration time (Fig. 1C) and exploration preferences during training did not show a treatment effect (discrimination index, Fig. 1D). As expected, in Overlapping a positive discrimination index was already seen in trial 5, due to the moving object-location during training. At test, there was a significant condition X treatment interaction (rmANOVA condition F_1,36_=5.6 p=0.023, treatment F_1,36_=1.2 p=0.28, interaction F_1,36_=5.1 p=0.031). CBD administration led to disrupted cumulative memory expression (Overlapping condition) but intact simple memory (Stable condition).

In sum, using a potentially translatable approach with oral administration of CBD, we showed a significant effect on behavior, where CBD led to intact simple but worse cumulative memory.

### CBD extends natural NonREM sleep

Our second aim was to characterize the effects of CBD on sleep and sleep oscillations. For this, new rats (n=4) were implanted with a wire-drive targeting the prelimbic cortex and hippocampus, which allowed us to acquire wake and sleep data in the task (25min) and sleep box (6h) for each study day (study design Fig.2A, histology Fig. S2). The recordings were divided into 5 min task and 45 min sleep-box bins (between each trial and 4 bins after the 5^th^ trial). Raw signal and spectrogram for one such 45min is presented in Fig.2A. The signal was manually scored for sleep stages NonREM, Intermediate and REM. Fitting to the idea that CBD promotes sleep, rats treated with CBD had more NonREM sleep but no changes in Intermediate and REM sleep nor the number of sleep stage transitions (Fig.2B-E). However, the effect on NonREM sleep was modest, it only became apparent in the last two sleep bins, when control rats showed a decline in amount of NonREM sleep. Of note, this is also the time period where the maximum level of CBD should have reached the brain.

**Fig.2.**
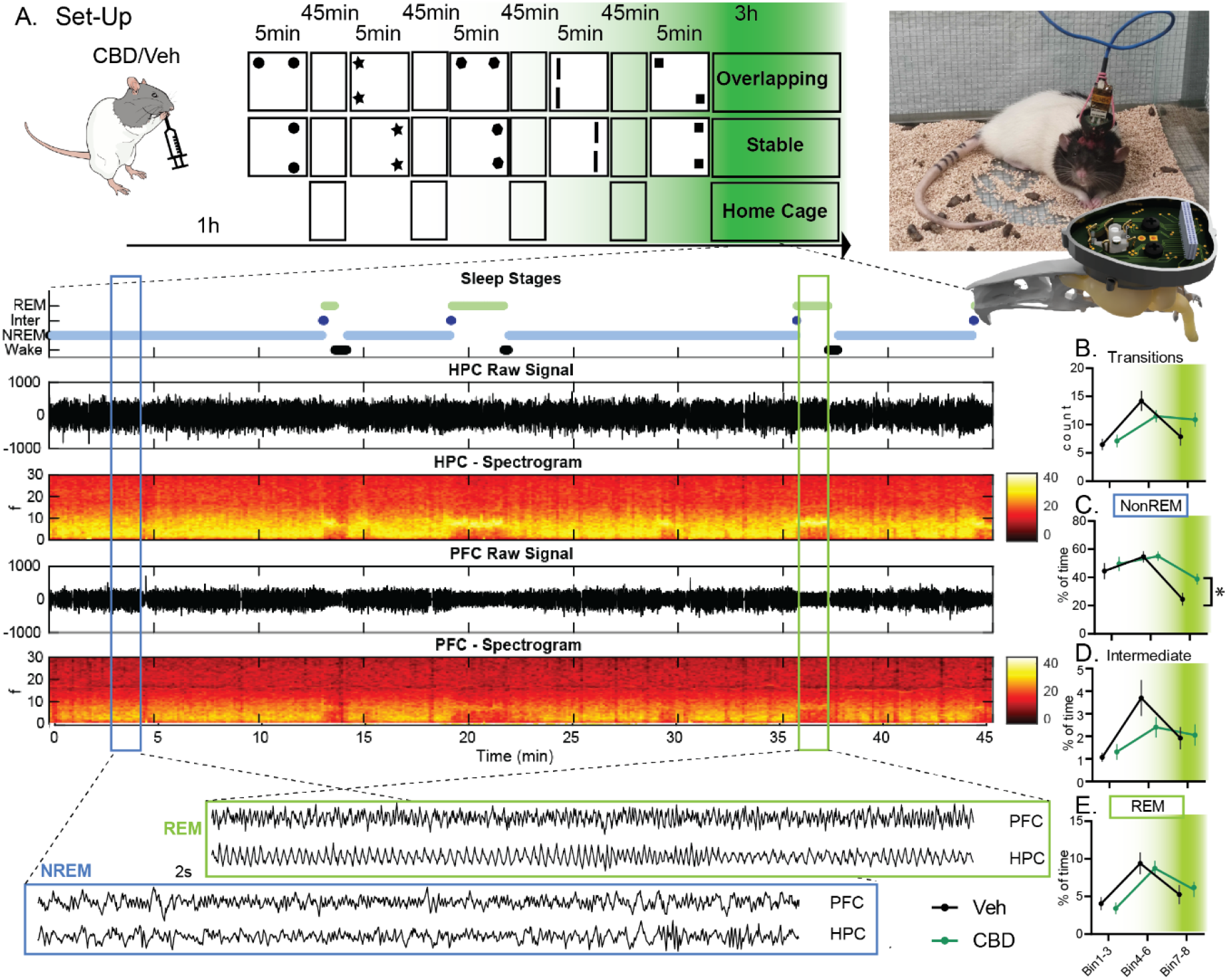
Electrophysiological experiments. A. Rats were implanted with a wire-drive (picture on the right). On study days they received CBD or Vehicle (Veh) orally and one hour later started the OS training as in Fig.1. In addition to the conditions Overlapping and Stable, animals also spent one study day always in the Home Cage during the usual training periods (circadian, no-learning control). Below manually scored sleep stages as well as raw signal and spectrograms arising from the hippocampus (HPC) and prelimbic cortex (PFC) are shown for an example 45min bin. Finally, for each NonREM (NREM) and REM state a 2s example of raw LFP is presented. Next separated for Bin 1-3, 4-6 and 7-8 the count of state-transition (B) as well as percent time spend in NonREM (C), Intermediate (D) and REM (E). Only NonREM showed a significant effect with more sleep in the final bins in rats treated with CBD. For A-E green shading implies expected effective CBD levels. *p<0.05

In sum, CBD led to more NonREM sleep at the end of the sleep period.

### CBD makes NonREM oscillations smaller

Next, we detected specific NonREM oscillations that are implied to be critical for memory consolidation [12]. Delta and spindle oscillations in the prelimbic cortex and ripple oscillations in the hippocampal CA1 area were detected during NonREM sleep periods. Deltas and ripples were much more numerous than spindles (Fig. 3A). To account for these differences, the counts of oscillations were normalized by their overall average and then, as for the sleep stages, separated for the different sleep bins. After CBD there were more deltas, spindles and ripples in the final bins (Fig. 3B); however, this was due to the increased amount of NonREM sleep, since there was no change in rates (Fig. S3). CBD did change the properties of each type of oscillation. Deltas and ripples became slower (decreased intrinsic frequency), while spindles became faster (Fig.3C). All three oscillations became smaller (Fig.3D).

**Fig.3.**
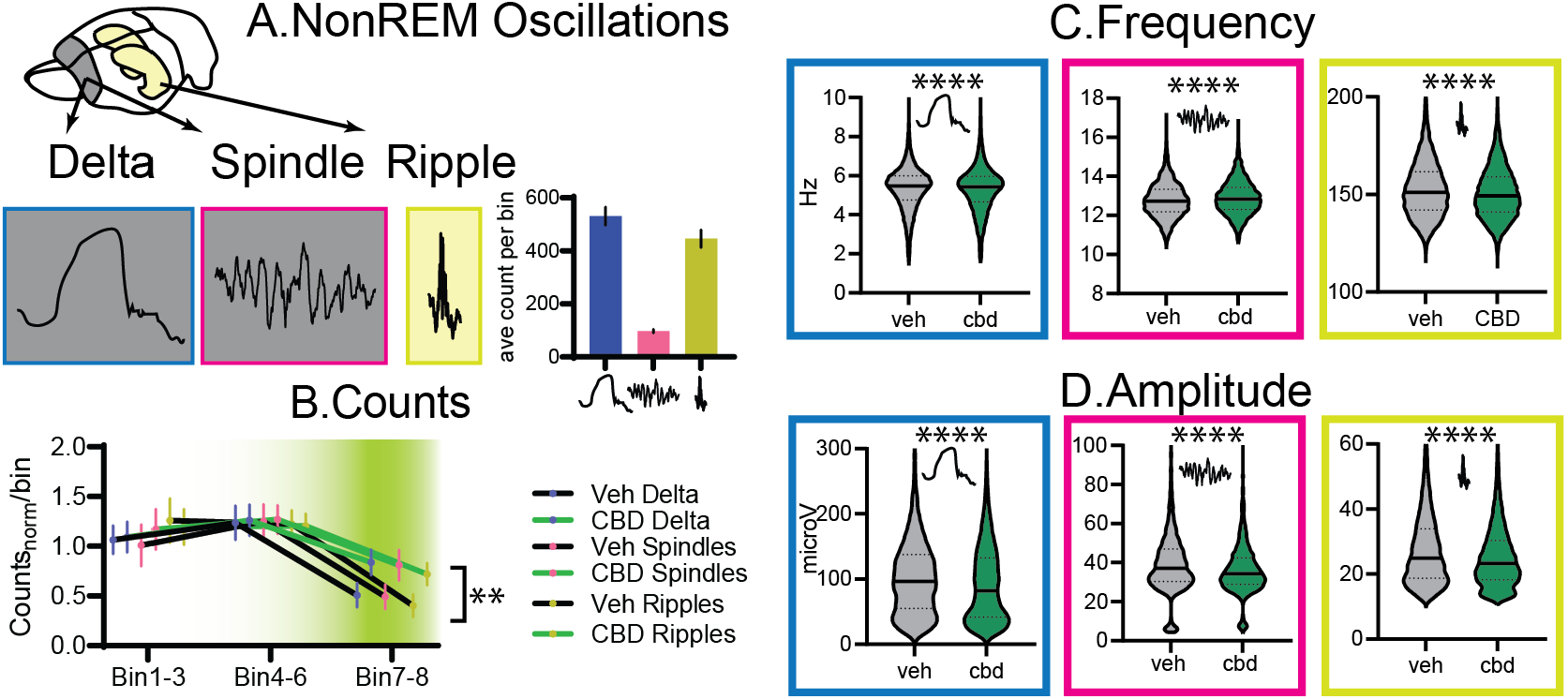
NonREM Oscillations. A. Cortical delta and spindles, and hippocampal ripples were detected. On the right, average counts per bin (types F_2,195_= 74.6 p<0.0001). B. Counts for all types normalized by their average. There was a significant treatment and time interaction (treat F_1,30_=5.2 p=0.029, bin F_2,40_= 30.9 p<0.001, treat X bin F_2, 60_=4.7 p=0.012 all other p>0.75). From left to right for delta, spindle and ripple the intrinsic frequency (C) and amplitude (D). Delta and ripples were slower and smaller, spindles faster and smaller (all K-S D, p<0.0001). ** p<0.01, **** p<0.0001

In sum, more NonREM sleep in CBD led to more NonREM oscillations at the end of the sleep period. However, CBD changed the frequencies and decreased the amplitudes of these oscillations.

### CBD decreases the number of long ripples

Memories are reactivated during sleep and these reactivations preferably take place during hippocampal ripples [41]. Recently, it has been shown that extending short ripples during a task led to better working memory, potentially because longer reactivations could be supported by the longer ripples [42]. Fittingly, in sleep it has been shown that longer ripples and ripple-trains support the reactivation of long memory-sequences on a 10m track [43]. Thus, long ripples during sleep could be especially important for memory consolidation of extended experience or more complex information. Therefore, next we split detected NonREM ripples into short (≤100ms) and long (>100ms) events (Fig.4A). Most events were short and these presented the same time-course as all ripples, with increases after CBD in the final sleep bins (Fig.4B). In contrast, long ripples occurred less already early in the day after CBD administration (Fig.4C). CBD decreased the amplitude of both short and long ripples (Fig.4D). Interestingly, while short ripples showed significant phase coupling in both VEH and CBD (both p<0.0001), long ripples were only coupled to the slow oscillation after VEH administration and not CBD (p<0.05, p=0.9, Fig.4E).

**Fig.4.**
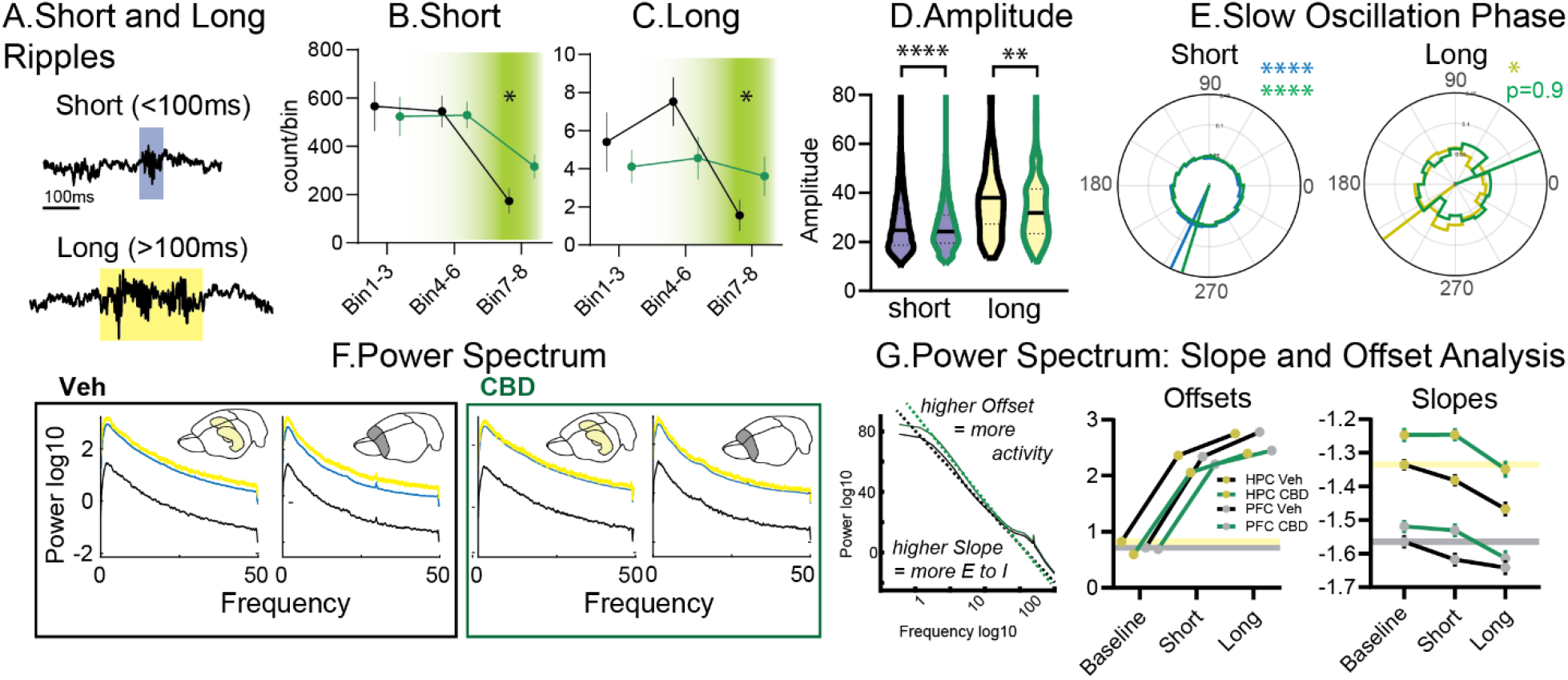
Short and Long Ripples. A. Examples. Counts for B. short and C. long ripples. D. Amplitudes. E. Slow oscillation phase, all short ripples were locked but for long ripples only in vehicles significant locking was seen. F. Power spectrum for vehicle and CBD, always left for hippocampus and right for prelimbic cortex. Of note, in Veh there was more power for long than short ripples. This was not seen in CBD. G. Slope and Offset Analysis from left to right: how slopes and offsets are calculated, offset and slopes for ripple types and random selected baseline. Lines indicate vehicle baseline levels. Offsets: events F_2, 3588_=1592 p<0.0001, treatment F_1, 3588_= 63.15 p<0.0001, events X treatment F_2, 3588_=4.5 p=0.011, brain area X treatment F_1, 3588_= 5.0 p=0.026. Slopes: event F_2, 3588_=24.6 p<0.0001, brain area F1, 3588=397.5 p<0.0001, treatment F1, 3588=47.3 p<0.0001, brain area X type F_1, 3588_= 6.2 p=0.0127 other p>0.27)

Plotting the hippocampal and cortical power spectrum during short and long ripples (in comparison to random selected baseline signal) revealed a frequency independent increase in power for all events to baseline. However, in vehicles long in comparison to short ripples had an even higher power, especially in the prelimbic cortex (Fig.4F). This was not evident in CBD. The power spectrum can be used to calculate E/I balances as well as overall activity levels. Specifically, from lines fitted to the double logged (power and frequency) power spectrum the offsets and slopes can be derived (Fig.4G). The offsets reflect the overall activity (higher offset → higher activity), while the slopes reflect the E/I balance (higher slope → higher E/I ratio). In vehicles, offsets (activity) in both brain areas increased from baseline to short and from short to long ripples, but the increase was less after CBD. Interestingly, slopes were generally higher in the hippocampus than in the cortex (higher E/I balance) and CBD increased these even more (especially in the hippocampus). After vehicle slopes decreased for ripple events (lower E/I balance). While this was similar in CBD, due to the overall higher baseline, slopes remained higher during ripples.

In sum, after CBD there were fewer long ripples and these were less locked to the slow oscillation phase. During especially long ripples, CBD showed less activity in the cortex. In general, CBD led to higher E/I ratio that could have led to the changes in ripple events.

### Only in Vehicle did cumulative-learning lead to more long ripples after delta waves

After establishing general effects of CBD on sleep, we next investigated the different memory training conditions in more detail. For this purpose, we analyzed NonREM oscillatory coupling, which is known to be a reliable learning marker. Hippocampal ripples interact with other NonREM oscillations. Couplings often increases after learning experiences and artificial enhancement of such couplings can strengthen weak memories [18]. However, many different types of Delta-Spindle-Ripple couplings have been reported and implicated to be important for memory consolidation. These couplings can be interactions between two oscillations such as delta followed by spindle [44], delta followed by ripple [45], ripple followed by delta [18], and spindles with a ripple in their troughs [46], but also three-oscillation interactions such as delta followed by spindle with a ripple in the trough [47], delta followed by ripple then spindle [48], ripple followed by delta and then spindle [18]. Until now, experiments tend to report only on one type of coupling, making it difficult to compare results and create a framework for potential different or common functions.

First, we focused on cortical events that can also be measured in human EEG experiments. Interestingly, the largest condition effect was seen for Delta-Spindle coupling (D-S), where only in the simple learning condition (Stable) there was an increase in D-S events that was not seen in single spindle events and was only weakly present for single delta events (Fig. 5A).

**Fig.5.**
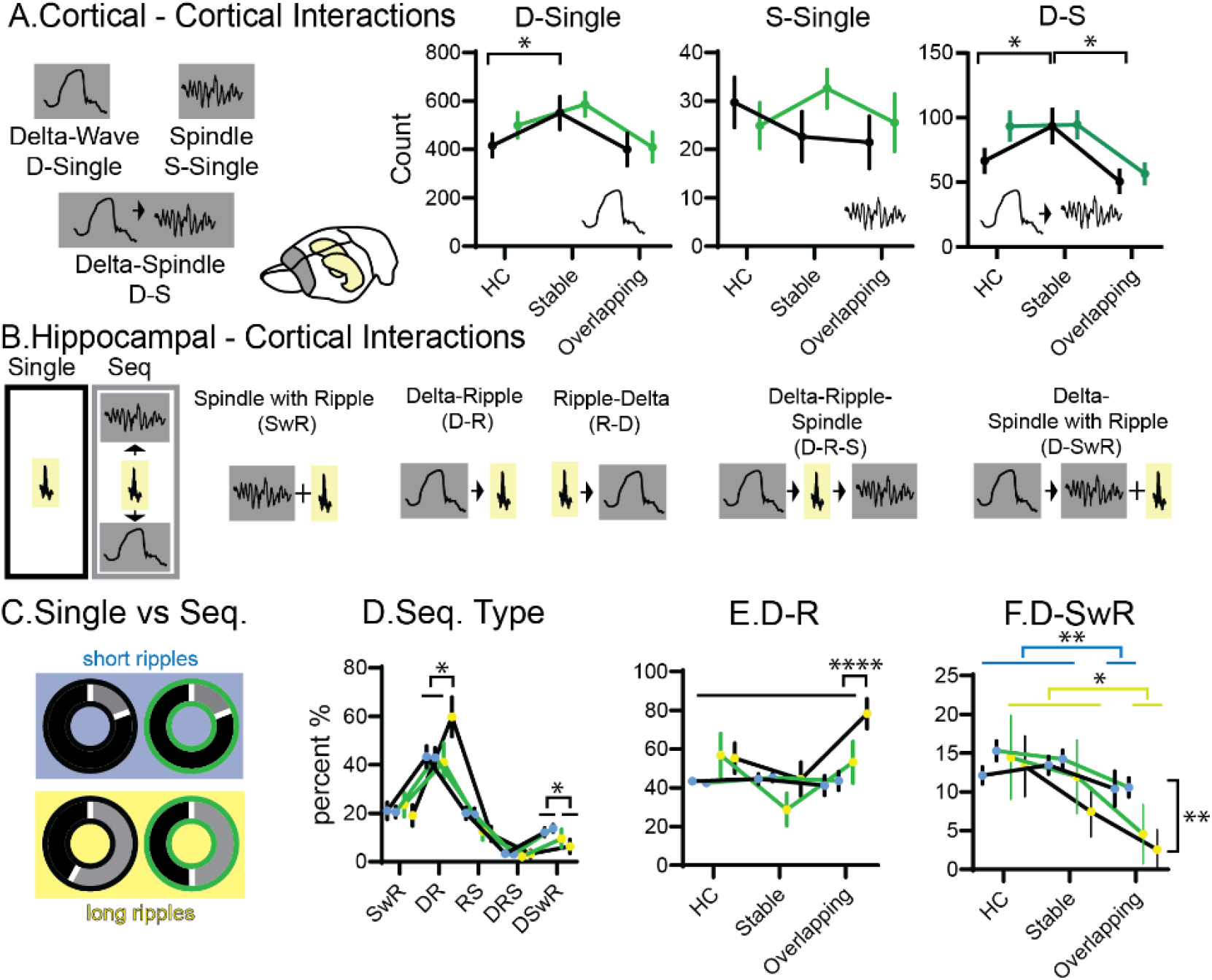
Oscillation interactions. A. Cortical interactions. Oscillations can be single delta (D) or spindle (S) waves, or coupled delta-spindle events (D-S). Counts of events shown for three different conditions home cage (HC), Stable and Overlapping (each rmANOVA with condition and treatment, D-single condition F_2,62_=3.9 p=0.026 other p>0.2, S-Single all p>0.13, D-S condition F_2,62_=6.2 p=0.0034, treatment F_2,62_=3.4 p=0.076, interaction p=0.29). B. shows different types of oscillatory coupling. C. Fraction of short (top blue background) and long (bottom yellow background) that are part of sequences (grey) or occur alone (black). Left vehicle (black edge), right CBD (green edge). D. Fraction of coupled long and short ripples spread over the different interaction types (sequence types F_4,160_=77 p<0.0001, sequence types X ripple types F_4,160_=2.3 p=0.06, other P>0.16). E. as in D fraction of events that are D-R but split for the different conditions (condition F_2,204_=5.2 p=0.006, ripple type F_1, 204_=7.2 p=0.0081, treatment F_1,204_=3.1 p=0.08, condition X ripple type F_2,204_=7.4 p=0.008, ripple type X treatment F_1,204_= 3.9 p=0.05, other p>0.2), F. same for D-SwR (condition F_2,204_= 6.5 p=0.0018, ripple type F_1,204_= 5.5 p=0.019 other p>0.18). Only in overlapping an increase was seen in vehicles but not CBD. * p<0.05, ****<0.0001

**Fig.6.**
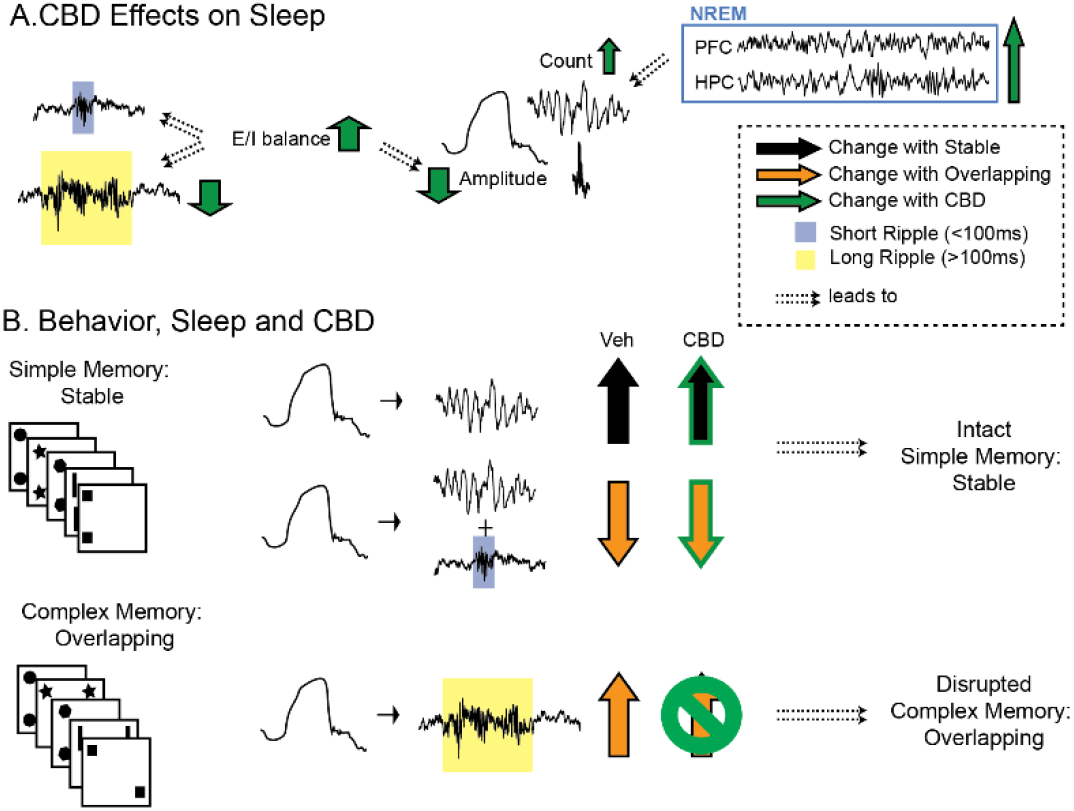
Summary. A. General CBD effects on sleep. CBD increases excitatory/inhibitory balance (E/I), which (potentially) leads to less long ripples and smaller NonREM oscillations. CBD also extends NonREM sleep and thus increases the number of NonREM oscillations at the end of the sleep period. B. Behavior, Sleep and CBD effects. After simple memory training (condition Stable) there is an increase in Delta-Spindle events in vehicle (VEH), which is unchanged in CBD. After complex memory training (condition Overlapping) there is a higher fraction of long-ripples following delta waves, but a smaller fraction of both short and long ripples during spindles following delta waves. The former but not latter effect is abolished with CBD, leading to disrupted complex memory consolidation in CBD.

Next, we analyzed different possible combinations of hippocampal-cortical interactions (Fig.5B): Delta-Ripple (D-R), Delta-Ripple-Spindle (D-R-S), Ripple-Delta (R-D), Spindle with Ripple (SwR) and Delta-Spindle with Ripple (D-SwR). Overall, long ripples were more likely to occur in one of the sequences than alone. In contrast, short ripples were much less likely to occur in a sequence. After CBD, these differences remained, however, now the fraction of long ripples in sequences decreased (Fig.5C). When splitting for sequence type, it became apparent that in vehicles, but not CBD, long ripples occurred more than short ripples as D-R events. In contrast, for both CBD and vehicle short ripples occurred more as D-SwR events than long ripples (Fig.5D). We delved deeper and could show that long ripple D-R events occurred less throughout the whole recording period in CBD. Finally, since long ripples should preferentially be associated to longer reactivations and these in turn would be associated to previous knowledge, we next split the recordings for the different conditions. The increase of long ripples occurring as D-R events shown in panel D was due to a selective increase of these events after the Overlapping in comparison to the Home Cage or simple learning control (Stable) for vehicles and not CBD (Fig. 5E). This condition-selective CBD effect mirrors the behavioral finding that only in Overlapping – testing complex, abstracted memory – animals showed worse performance with CBD. Interestingly, D-SwR events were less in overlapping for both short and long ripples (Fig. 5F). In sum, long ripples are more likely to occur in sequences than short ripples. After learning a complex memory (Overlapping) a selective increase in the fraction of long ripples that occured after delta waves was seen and accompanied by a decrease in D-SwR events. The increase in D-R was absent in CBD and potentially led to the behavioral deficit in this memory condition. After the simple memory condition (Stable) and increase in D-S events was seen.

## Discussion

We investigated the effects of oral CBD intake on memory consolidation and sleep in rats. We could show that CBD extended natural sleep but also led to a higher Excitatory/Inhibitory (E/I) balance, smaller amplitude NonREM oscillations and less long ripples in NonREM sleep. Only after vehicle treatment, complex learning resulted in an increased fraction of long-ripples following delta-waves, a potential consolidation mechanism. In CBD, this condition-specific increase in oscillation-coupling was not seen and the rats showed deficits in complex memory expression the next day. In contrast, simple learning led to an increase in delta-spindle coupling events in both vehicle and CBD, and following both treatments intact memory expression for simple memory was seen the next day. Thus, while we can confirm the beneficial effect of CBD on sleep length, this effect was modest and accompanied by changes in E/I balance and NonREM oscillations that led to a deficit in complex memory consolidation.

### CBD and E/I balance

CB1-receptors are found on the presynaptic terminals of interneurons, thus a main neuronal effect of systemically-administrated CBD, should be a change in E/I ratio. Due to this, CBD is currently being handled as a novel therapeutic agent restoring E/I balance in neural models of epilepsy and autism [4, 5] in addition to being used increasingly as sleep medication.

We know that activation of the CB1-receptors with synthetic agonists inhibits hippocampal GABAergic neurotransmission and dysregulates the distribution of interneurons [49, 50]. What remains unclear is in which direction CBD administration changes the E/I ratio. Depending on the dose, CBD can have different effects as functional agonist or as antagonist of the CB1-receptor. In low concentrations CBD acts directly as an antagonist to CB1 receptors [51] but it can in other doses also function as an antagonist of CB1-receptor-agonists [24]. Thus, actions of CBD are complex and depending on the dose, it can increase or decrease inhibition. This may also explain different effects on sleep depending on dosages per study. High doses increased sleep durations [6] and the opposite effect was seen with low doses [7, 52]. But complexity does not stop there, looking at studies done in rodents, different endocannabinoids seem to be active at different phases of sleep, underlying complex interactions with NMDA and AMPA receptors [53]. The effects also seem to be dependent on period of administration – acute or chronic.

Here, we aimed to model an acute, high-dose intake, as would be recommended in humans to promote sleep. We could replicate the finding that high-dose administration increased total sleep time, even though the effect was modest and was seen as an extension of the NonREM sleep period at the end of the day. By analyzing the slopes of the power spectrum, we could determine how CBD changed E/I ratio. Interestingly, during NonREM sleep periods CBD led to an increase in E/I ratio, likely by decreasing inhibition. This was visible in the prefrontal cortex but was even more evident in the hippocampus. Perhaps the change in E/I balance contributed to the extension of sleep. While a direct relationship between E/I balance and sleep is not yet known, we do know that interneuron activity is necessary for most sleep oscillations as we will expand on next.

### CBD, E/I balance and NonREM oscillations

NonREM sleep is dominated by the transition of up- and down-states (states of general neuronal activity and silence, respectively). Large down-states can induce a delta-wave in the LFP signal, which we can detect. Interestingly, two major classes of interneurons, the parvalbumin and the somatostatin positive cells, tightly control both up-to-down and down-to-up state transitions [54]. The activity of these interneurons is needed to initiate and maintain the down-state [54]. Further, sleep spindles are associated with local, cortical parvalbumin interneuron activity measured with calcium imaging [55], which again is potentially necessary to maintain this oscillation. Finally, hippocampal ripples are initiated by excitatory, pyramidal cell activity but then maintained by activity of interneurons [56]. Thus, any effects of CBD on these sleep oscillations, are likely due to its effect on inhibition. Already previously it had been shown that administration of anandamide (CB1-agonist) disrupts ripples in the hippocampus [57]. This disruption effect was abolished by administration of CB1-receptor antagonist, evidence that CB1-receptor activation was the causal link. On similar lines, injection of CB1-receptor agonists in hippocampal cultures also disrupted ripple events [58].

Here, we could show that CBD affects ripples and other sleep oscillations as well. Deltas and ripples became slower, spindles faster. Importantly, all three oscillations became smaller in amplitude, likely due to decreased entrainment of excitatory neurons by the now attenuated interneurons. E/I balance was changed more in the hippocampus than in the cortex and we also saw a more complex picture of CBD interactions with hippocampal ripples. During ripples normally E/I balance decreases, this decrease was seen less in CBD and, due to higher E/I baseline levels in CBD, event E/I balance remained higher than the baseline of vehicles. A specific E/I balance is needed for ripple maintenance, correspondingly we saw fewer long ripples in CBD. Further, those long ripples that remained in CBD showed smaller cortical, wide-band responses in the spectrogram. Critically, in CBD the condition-specific increase in coupling of large down-states to long ripples (D-R sequences) after complex learning was absent. Likely leading to the memory deficit seen after CBD treatment in this condition. In contrast, while there were E/I differences in the cortex, these were smaller than the differences in the hippocampus. Correspondingly, there was no CBD effect in the count of cortical oscillation-coupling events (delta-spindle), that were associated to simple-learning, and CBD also did not lead to deficits in this memory condition.

In sum, by changing excitatory and inhibitory neuronal balance, CBD led to changes in hippocampal-cortical oscillations and coupling resulting in deficits consolidating complex learning events. In contrast, within-cortex coupling remained intact, which likely was why CBD treated rats did not show deficits in simple learning.

### Oscillation-Coupling for Memory Consolidation

At this point we do know that ripples, spindles and delta-waves in sleep are important for memory consolidation. Further, for some of their functions these oscillations are coupled to each other, as reported by many researchers. Surprisingly, no two studies seem to be in agreement, which coupling is the one to look for. Reported couplings can be interactions between two oscillations such as delta followed by spindle [44], delta followed by ripple [45], ripple followed by delta [18], and spindles with a ripple in their troughs [46], but also three-oscillation interactions such as delta followed by spindle with a ripple in the trough [47], delta followed by ripple then spindle [48], ripple followed by delta and then spindle [18]. Experiments tend to report only on one type of oscillation, and different types of memories as well as different coupling events are never directly compared.

Here, we systematically investigated all these couplings directly comparing a non-learning control (HC) to both a simple and complex memory condition (Stable and Overlapping, respectively). First off, the simple memory condition (Stable), which has the different objects placed at the same two locations throughout training and at test one new location is used, is similar to classic reference memory such as the watermaze [59, 60]. Since the different trials have the same configurations, the animal can use both a cumulative memory across trials as well as a recency memory of the last trial to still perform above chance. In contrast, in the Overlapping condition the objects will be presented in different locations from trial to trial, the trial-by-trial novelty should encourage the animal to consolidate these experiences during sleep and abstract the underlying cumulative statistical distributions. In Overlapping this distribution is skewed, such that one location will always contain an object. Thus, we can test for the behavioural expression of the cumulative memory, by presenting the same spatial configuration at test as it was presented during the final trial. Now, only if the animal extracted the cumulative statistics during all training trials, will there be a preference for the less often shown location at test.

By contrasting these different conditions, we can extrapolate which effects are driven by which type of learning or experience. Changes induced by all OS conditions in comparison the to Home Cage, should represent more general experience-dependent effects enabling simple memory consolidation or homeostasis. Changes seen only after Stable condition, would represent consolidation of simple memories, already reinforced during training. In contrast, changes seen specifically after Overlapping should represent semantic-like memory consolidation and the comparison and integration of new and old information.

Interestingly, here we can show in vehicles that Stable led to more Delta-Spindle events. In contrast, in Overlapping the fraction of long-ripples following a delta-wave increased while the fraction of ripples occurring during a spindle after a delta wave (D-SwR) decreased. We had already previously observed that the increase of delta and spindles and their coupling was linked to simple experiences [14], and this coupling is most often reported by researchers investigating human subjects with simple, associative memory paradigms such as paired-associative word-list or reinforced spatial learning [47, 61]. This fits to our current results, where delta-spindle coupling increased after our simple memory condition (Stable). In contrast, ripples following delta are reported, for example, in experiments focused on prelimbic reactivations when animals learn complex rules [45]. In our results, this coupling increased specifically after the complex learning conditions. It would be tempting to speculate that Delta-Spindle and Spindle with ripple coupling would facilitate simple learning while Delta-Ripple would correspond to consolidation of complex memories relying during consolidation on the comparison to other previously encoded experiences. Our current results support this, since after CBD the condition-specific Delta-Ripple increase was absent but Delta-Spindle coupling remained the same, and correspondingly CBD only induced a memory-deficit in the complex and not simple memory condition.

In conclusion, by combining oral CBD administration with the Object Space Task and electrophysiological recordings, we could show that CBD leads to 1) extension of NonREM sleep, 2) increased E/I balance, 3) smaller NonREM oscillations, and 4) fewer long ripples and less delta-long ripple coupling. The latter leads to 5) deficits in complex-memory consolidation but simple memory remains intact after CBD.

## Acknowledgments

We would like to thank Linda Tomy for running some analysis and Eloise Kuijer, Vera Mascarenhas Pombeiro Duarte Silva, Nienke Flipsen for running the behavioral experiments.

## Contributions

AS supervised and performed experiments as well as data analysis, AAZ supervised and performed data analysis, KA and PO analyzed the electrophysiology data, AA, JvdM, INL helped with experiments and supervision, AR supported electrophysiology data analysis, LG designed and supervised the project. AS, LG and AAZ wrote the first draft of the manuscript.

---

**Fig. S1:**
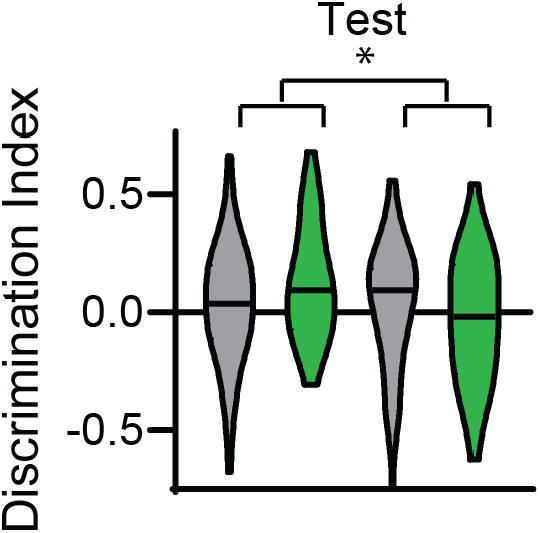
Shown is the test data for all rats. The interaction CBD and Condition remained significant.

**Fig. S2:**
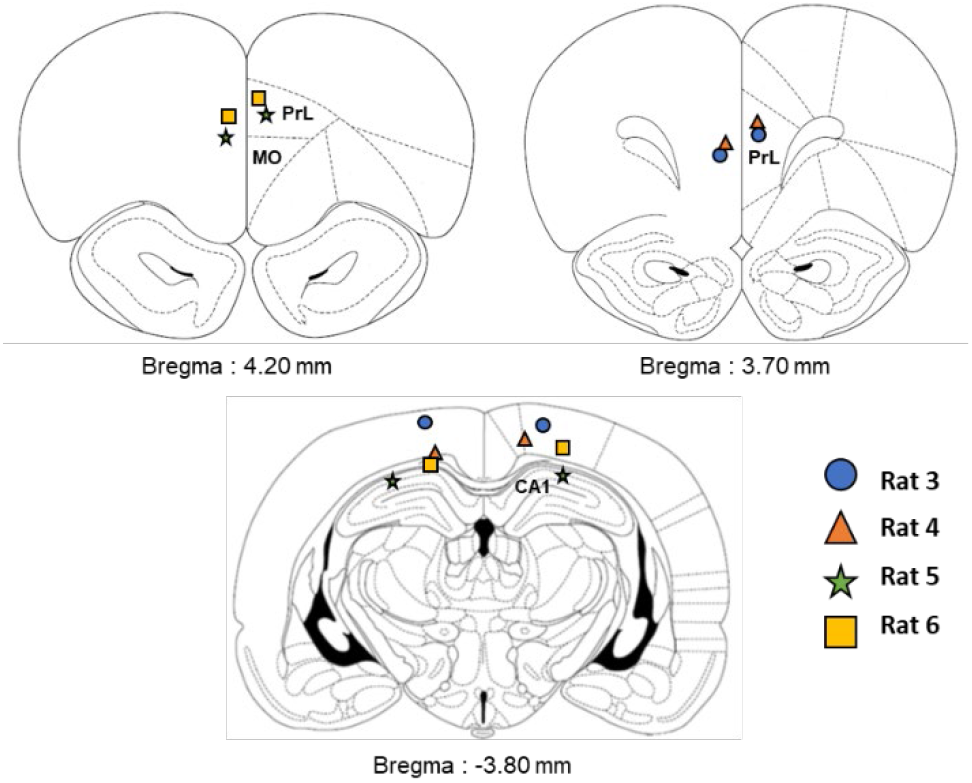
Placement of electrodes after histological confirmation. Rat 3 did not reach the hippocampus and was only included for analysis of sleep stages and cortical events. All the other rats had at least one electrode with ripples.

**Fig. S3:**
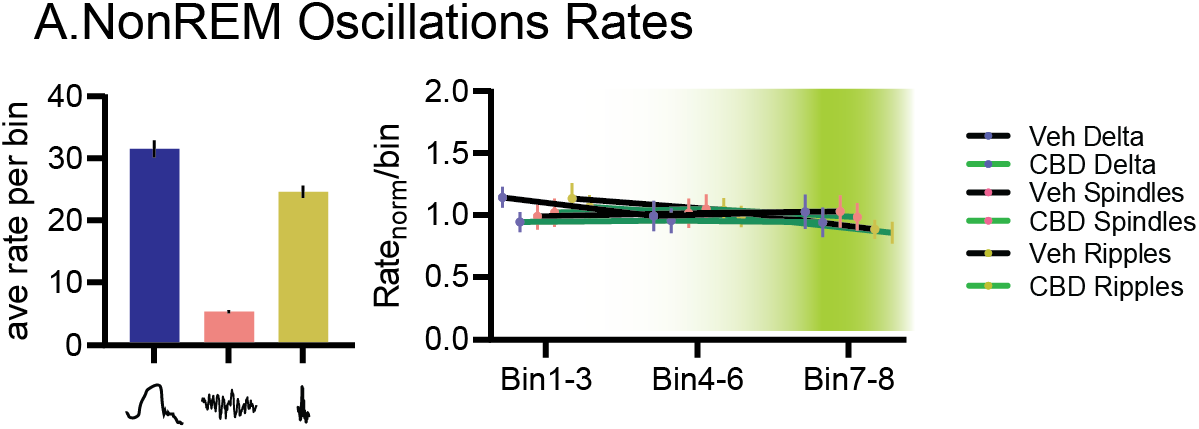
Rates of NonREM oscillations

